# Managed honey bee colony losses and causes during the active beekeeping season 2022/2023 in nine Sub-Saharan African countries

**DOI:** 10.1101/2024.04.30.591982

**Authors:** Beatrice T. Nganso, Workneh Ayalew, Abebe J. Wubie, Freweini Assefa, Lulseged Belayhun, Nelly N. Ndungu, Daniel Toroitich, Z. Ngalo Otieno-Ayayo, Mbatha B. Wambua, Yudah O. Oyieyo, Ntirenganya Elie, Rachidatou Sikirou, Souradji B. Idrissou, Willy Mwiza, S. Turner, Bridget O. Bobadoye, Sidonie T. Fameni, Sayemie Gaboe, Mawufe K. Agbodzavu, Patrick Mafwila, Geraud C. Tasse Taboue, Kimathi Emily, Tonnang Z.E. Henri, Saliou Niassy, Simplice N. Fonkou, Christian W. W. Pirk, Alison Gray, Robert Brodschneider, Victoria Soroker, Sevgan Subramanian

## Abstract

This study reports for the first-time managed honey bee colony loss rates and associated risk factors during the active beekeeping season 2022/2023 in nine Sub-Saharan African countries, namely Kenya, Ethiopia, Rwanda, Uganda, Benin, Liberia, Nigeria, Cameroon and Democratic Republic of the Congo. The sustainability of bee swarm catches as a main honey bee colony source tool for operation expansion by African beekeepers was also evaluated in Kenya and Ethiopia. In this survey, the 1,786 interviewed beekeepers across these countries collectively managing 41,761 colonies registered an overall loss rate of 21.3%, which varied significantly among countries (from 9.7 to 45.3%) and hive types (from 10.6% in hives with movable frames to 17.9% in frameless hives). The perceived causes of losses in order of significance were issues beyond the beekeeper’s control (mainly theft, drought, and bushfire), absconding and pests (mainly wax moth, small and large hive beetles, ants and *Varroa destructor* mite), but this pattern varied greatly across countries. Among the management practices and characteristics, migratory operations and professional beekeepers experienced lower losses than stationary operations and semi-professionals and hobby beekeepers. Insights into the number of bee swarms caught revealed significant decreases in swarm availability over the past three years in Kenya. The opposite situation was observed in some regions of Ethiopia. These trends require further investigation. Overall, this comprehensive survey sheds light on the complexities and challenges beekeepers faced in Sub-Saharan Africa, pointing to the need for targeted interventions and sustained research to support the resilience and growth of the apicultural sector.

## Introduction

Over the past 17 years, countries in the Northern hemisphere, particularly the United States, Europe and Canada, have consistently reported significant winter, summer and/or annual colony losses of the honey bee, *Apis mellifera* L. [1–7]. These losses are of economic significance to the apicultural and agricultural sectors as well as to the environment [8–10]. Several factors, which sometimes act synergistically, are responsible for these honey bee colony losses [2,3,11–13]. These include mainly invasive pests (particularly *Varroa destructor*), pathogens (particularly viruses associated with *Varroa* mite), issues beyond the beekeepers’ control (e.g. pesticides, extreme weather conditions and natural disasters like flooding, fire or vulnerability to theft) and management practices (e.g. hive migration and queen replacement) [2,3,11–13]. The impact of these stressors on colony mortality varies across countries and seasons [7,14].

As opposed to the situation in countries of the Northern hemisphere, reports indicate that colony losses in the Southern hemisphere (e.g. Africa, South America and Australia) have not been severe [6,7,15,16]. However, this might not be entirely true as there is great paucity of long-term spatiotemporal surveys that document managed honey bee colony losses and their causes in this part of the world [6,17]. For example, a few large-scale and spatiotemporal studies have reported above 30% colony losses in a number of Latin American countries [18,19], while below 25% losses were reported in Australia (PHA, 2019) and New Zealand [14]. In the African continent, the quantification of colony losses and causes has been restricted to a few countries in the south and north. In South Africa, high total colony loss rates of 29.6% and 46.2% in 2009-2010 and 2010-2011, respectively, were reported during the active beekeeping season from 1^st^ September 2009/2010 to 1^st^ April 2010/2011 [20]. These losses were more severe in migratory than stationary beekeeping operations, and small hive beetles, *Varroa* mites, absconding and chalkbrood disease were identified as key factors responsible for these losses [20]. In North Africa, Egypt experienced winter loss rates of 35.5% in 2011-2012 and 38.8% in 2012-2013, primarily due to *Vespa orientalis*, starvation, *Varroa* mite and poor quality of queens [21]. These losses were generally higher than the commonly estimated winter loss rate of 16% in European and some non-European countries [12]. However, in recent years, overall winter colony loss levels were somewhat lower at 24.3% in 2019-2020 for Egypt and 9.8-12.2% in 2017-2018, 2018-2019 and 2019-2020 for Algeria, with queen problems and natural disasters cited as the main factors [4,5,14]. These spatially limited studies are in vast contrast to the scale of beekeeping across Africa, highlighting the huge gaps in reporting colony loss data in most parts of the continent.

Apiculture is practiced in a different way across Africa compared to most parts of the world [22]. Of the many millions of colonies spanning eleven endemic honey bee subspecies occurring in this continent [23,24], only a small fraction are managed bees [25]. Most African beekeepers predominantly rely on capturing bee swarms to sustain and expand their apiaries during the active beekeeping season. This season is characterized by swarming, migration of colonies, absconding and honey harvest, with little routine management interventions at the colony level [26–30]. Thus, the factors driving managed honey bee colony losses in Africa are likely to be different from those reported in the Western world, influenced not just by genetic and environmental factors but also by distinct management practices, cultural, and socio-economic factors. In fact, many African beekeepers utilize various hive types ranging from movable to frameless structures [29,30]. Typically, they do not select for traits such as low absconding/swarming or defensiveness, despite these behaviours being more pronounced in African subspecies compared to their European counterparts [23,31]. The adoption of routine apiary management practices such as hive inspection, requeening, pest control, and provision of water and/or supplementary feeding is minimal among African beekeepers [20,29]. Additionally, a lack of education and experience in good colony management and harvesting techniques tailored to the hive types used can exacerbate the absconding rate and/or colony mortality [27,29]. Furthermore, minor apicultural pests such as wax moths, large hive beetles and ants reportedly cause up to 50% loss of managed honey bee colonies in certain parts of Africa [29,32]. Meanwhile, the impact of the ectoparasitic *Varroa* mite and its associated viruses is poorly documented in most African countries, and their significance in causing colony losses in the apicultural industry remains poorly studied. In countries like South Africa [33,34], Kenya [35–37] and Ethiopia [39], the impact of *Varroa* and its associated viruses on managed honey bee colonies is considered insignificant.

In this study, we attempted for the first time to quantify and compare loss rates of managed honey bee colonies during the active beekeeping season in nine countries in Sub-Saharan Africa. Additionally, we explored potential risk factors contributing to these losses. We further compared the colony loss rates based on several criteria: the country of operation, the type of beekeeping operation (professional, semi-professional, traditional or hobbyist), training in best beekeeping practices, beekeeping activity (migration versus non-migration), the types of hives used by the beekeepers, and colony management practices (provision of supplementary feeds and/or water at the onset of swarming to enhance colony establishment).

## Methods

### The survey

In this study, the questionnaire used was adapted from the 2023 COLOSS colony winter loss monitoring questionnaire to suit the specific conditions of beekeeping in Africa. The original COLOSS questionnaire refers to the non-active beekeeping season (winter) and not the active beekeeping season in Africa. The English version of the adapted questionnaire was distributed to national coordinators who spearheaded the survey (as in the COLOSS survey; [12]) from Eastern (Kenya, Ethiopia, Rwanda, and Uganda), West (Benin, Liberia, and Nigeria) and Central (Cameroon and Democratic Republic of the Congo) Africa. These national coordinators were selected from among the local advisors in research organizations and/or private sectors. They collected responses from beekeepers primarily through in-person interviews, seeking verbal consent from the respondents prior to data recording. Additionally, the coordinators from Kenya, Rwanda and Benin handed out questionnaires to beekeepers during workshops and meetings and follow-ups were carried out by phone calls to complete the data collection process. When necessary, the questionnaire was translated into French and/or local languages during data collection.

Our adapted questionnaire refers to the period from 1^st^ September 2022 to 30^th^ June 2023. This period captured the active beekeeping season and honey bee activity in the surveyed African countries. Some of the surveyed regions experience more than one swarming season during this beekeeping season. Accordingly, the total number of colonies managed changes over the season, given that most African beekeepers set out empty hives on trees and other standing structures to catch migrating honey bee swarms to expand their apiary size. As in the COLOSS questionnaire, the questions were categorized as mandatory (marked with an asterisk) or optional (without an asterisk) (Table 1). In the mandatory questions (Q), we asked about the disease symptoms and the loss rate of production colonies, defined as colonies with a queen that can provide a honey harvest during the active beekeeping season. We also evaluated the effect of beekeeping experience, training in best beekeeping practices, beekeeping activity, colony management and hive types on colony loss rates.

**Table 1.**
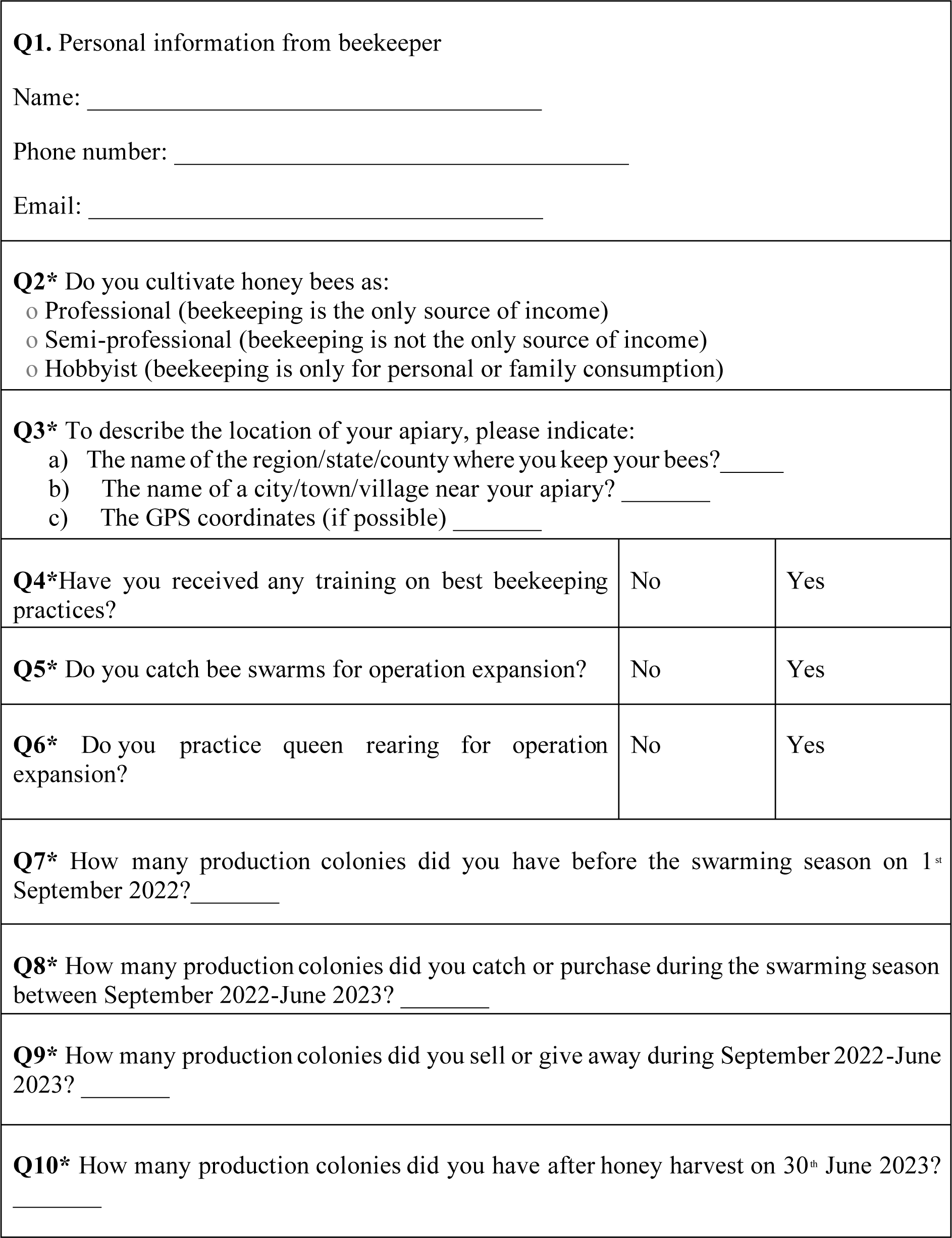
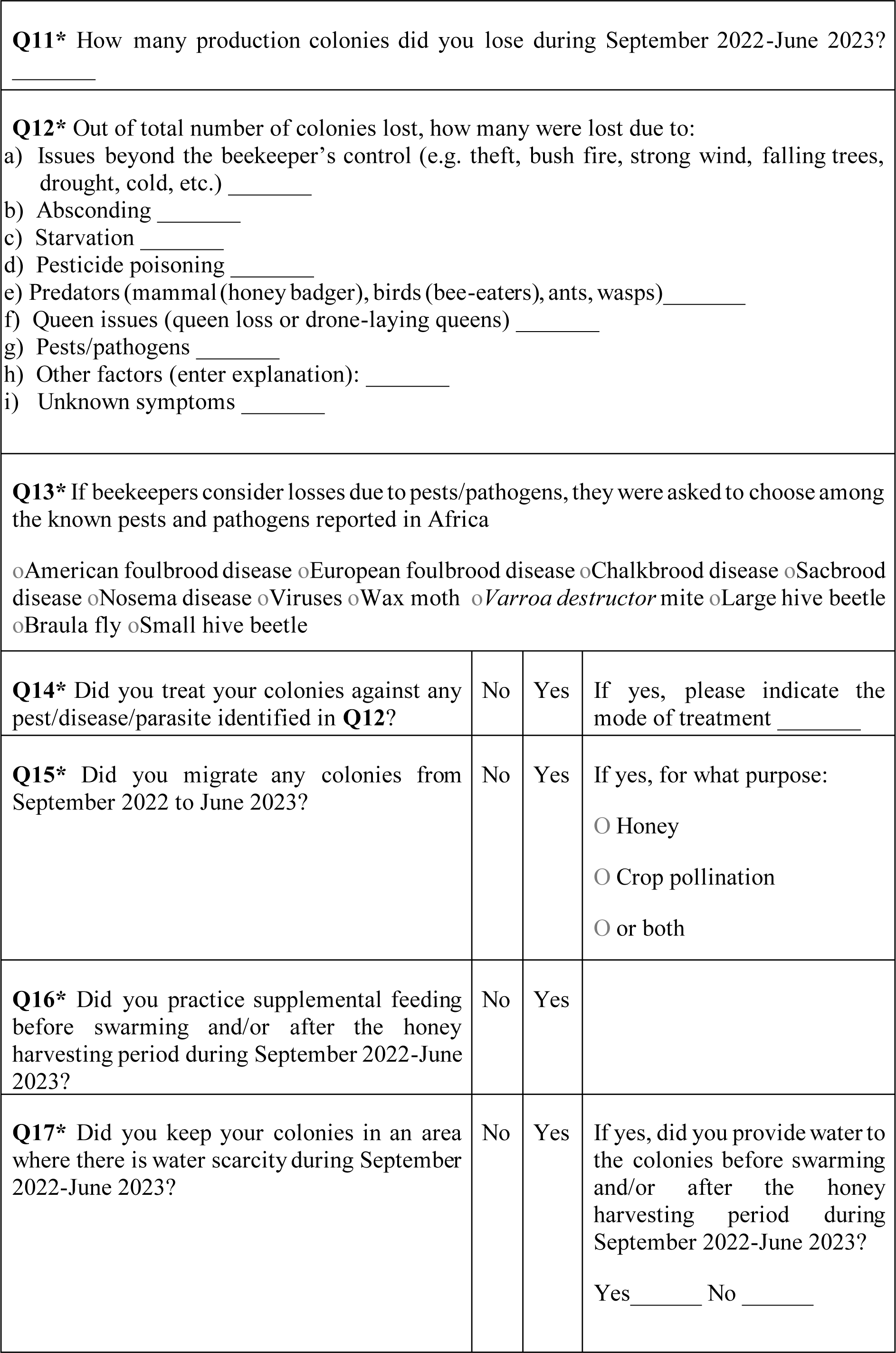
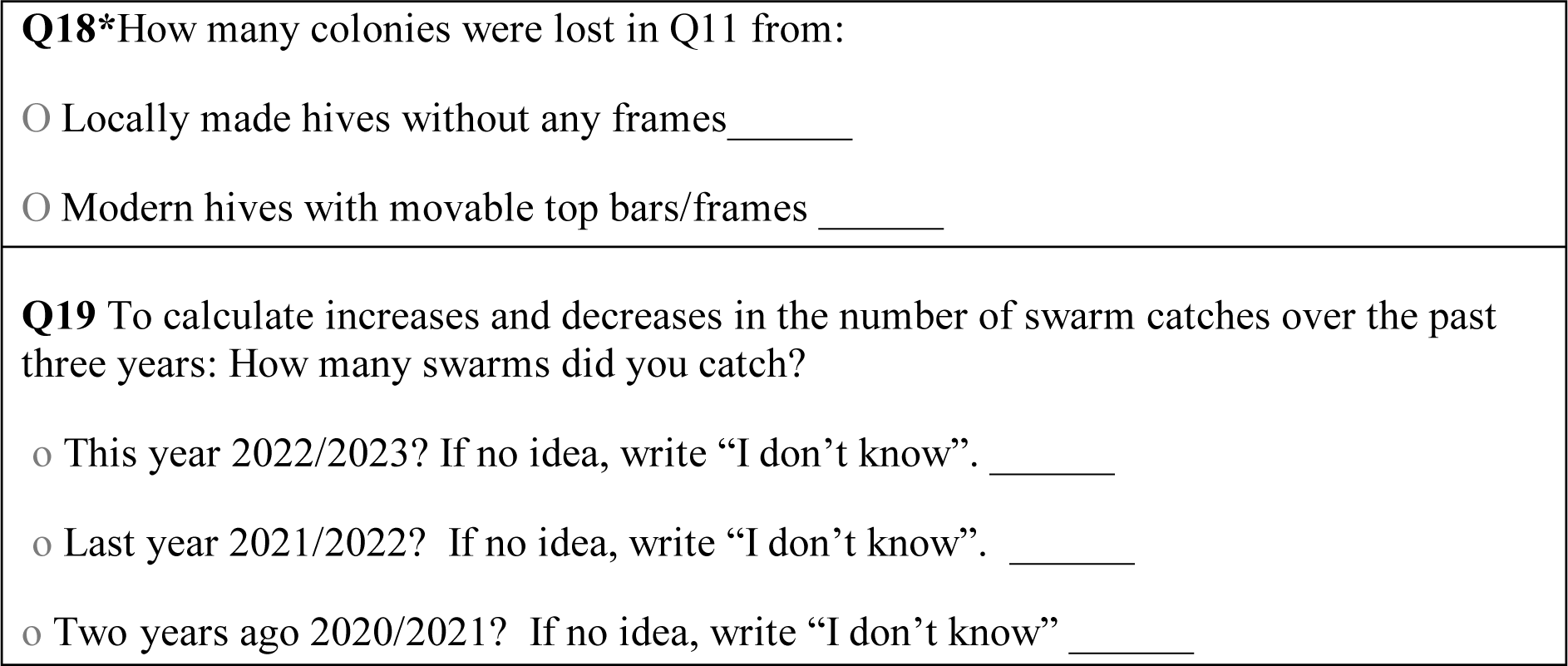
The questions asked in the survey for 2022/2023; the asterisks indicate mandatory questions.

The national coordinators calculated the colony loss rate in **Q11** during the interview section using the following formula: [(**Q7** + **Q8**) – (**Q9** + **Q10**)]. Also, for losses attributed to pathogens, care was taken by the national coordinators to explain the symptoms of viral diseases, microsporidia, and bacterial diseases to the beekeepers, as most of them are not knowledgeable about this topic. The **Q18** was only added to the questionnaire in June 2023. Fortunately, a few national coordinators were able to revisit the beekeepers to gather this information.

### Statistical analyses

All statistical analyses were conducted in the R-software version 4.3.2 [40]. All the raw data were checked for consistency before estimation of the total loss rate per country, as follows: (a) colonies at the start of swarming should not be missing and must be greater than zero; (b) colonies lost should not be missing and must be greater than or equal to zero; and (c) the total loss rate attributed to various risk factors should not exceed the total colony loss rate per country. Only valid data were used to estimate the overall loss rate for all the nine surveyed countries, to compute the 95% confidence interval (CI) for the overall loss rate, and to determine how country, the beekeeper’s profession, hive types and beekeeping activity influence the overall loss rate, the quasi-binomial family of generalized linear models (GLMs) were fitted using the R codes available in the Standard Survey Methods chapter of the COLOSS BEEBOOK [41] and the significance level alpha was set at 0.05. We also estimated the effect of the individual factors mentioned above within countries such as Ethiopia, Kenya, and Benin, where substantial data were available to report reliable results. The Poisson GLM fitted with a log link function was also used to analyse changes in the number of bee swarms caught over the past two years in Ethiopia and Kenya. This statistical approach was also used to swarm catches between Ethiopia and Kenya as well as among regions within Ethiopia compare over time. Means and t-intervals are quoted for the numbers of colonies managed in different countries and for various subgroups of beekeepers.

## Results

### Loss rates of managed honey bee colonies across the countries and within Ethiopia, Kenya and Benin

A total of 1,786 beekeepers from the nine surveyed African countries provided valid loss data for 41,761 honey bee colonies managed over the active beekeeping season (Table S1, Figure 1). This number of respondents represents less than one percent of the total number of beekeepers in the nine surveyed countries. In this survey, 95.3% of the respondents depended on swarm catches for operation expansion, whereas only 4.7% of them practice queen rearing for the same purpose.

**Figure 1.**
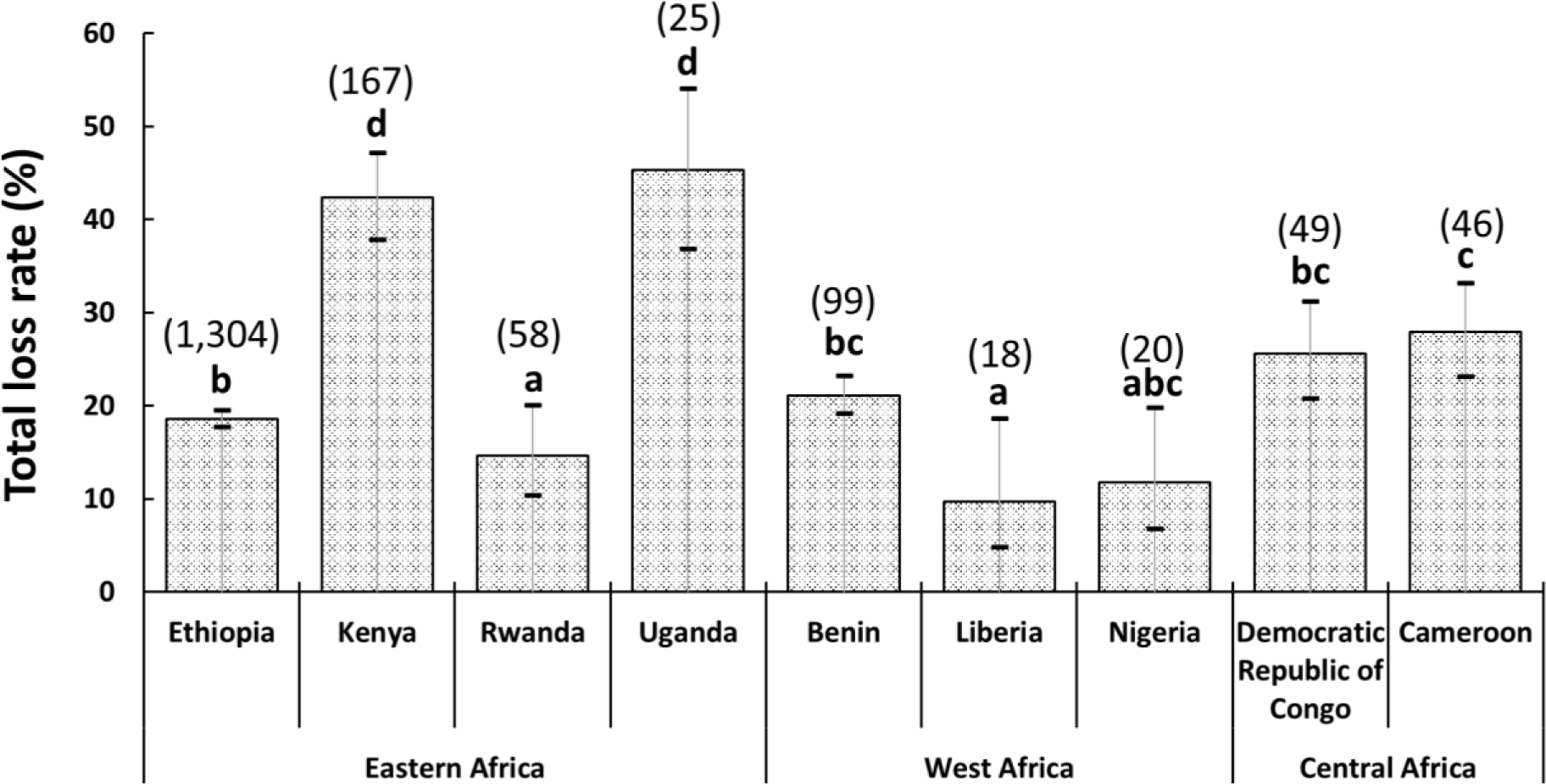
A bar graph showing the overall colony loss rate and 95% confidence interval for the 1,786 beekeepers interviewed across the nine Sub-Saharan African countries. Different letters above bars indicate significant differences among the countries when compared using the quasibinomial GLM with “country” as an explanatory factor in van der Zee et al. [41], *p* < 0.05. The numbers in brackets above bars indicate the total number of interviewed beekeepers who provided valid data for each country.

Following the honey harvest, the 1,786 beekeepers ended up with 32,861 colonies out of 41,761. Thus, the total number of colonies that were lost was 8,900. This represents a total loss rate of 21.3% (95% CI: 20.4-22.2%). The loss rate varied significantly among the surveyed countries (quasibinomial GLM: F = 41.5, df = 8, *p* < 0.001) (Figure 1). In fact, Uganda and Kenya registered the highest loss rates, whereas Liberia followed by Nigeria and then Rwanda registered the lowest loss rates. Loss rates in Ethiopia and Cameroon were intermediate and significantly different from each other and from Liberia and Rwanda, while loss rates in Benin and Congo were not significantly different to those in Ethiopia and Cameroon.

The total loss rate also showed significant differences among regions within Ethiopia (quasibinomial GLM: F = 19.7, df = 2, *p* < 0.001) (Figure 2A) and Kenya (GLM. quasibinomial: F = 7.3, df = 3, *p* < 0.001) (Figure 2B), but not among those in Benin (quasibinomial GLM: F = 3.02, df = 2, *p* = 0.05) (Figure 2C). In Ethiopia, beekeepers from the Southern Nations, Nationalities, and Peoples’ (SNNP) region registered lower losses than those from the Amhara and Oromia regions, while in Kenya, those from the Coastal region registered higher losses than those from the other three regions.

**Figure 2.**
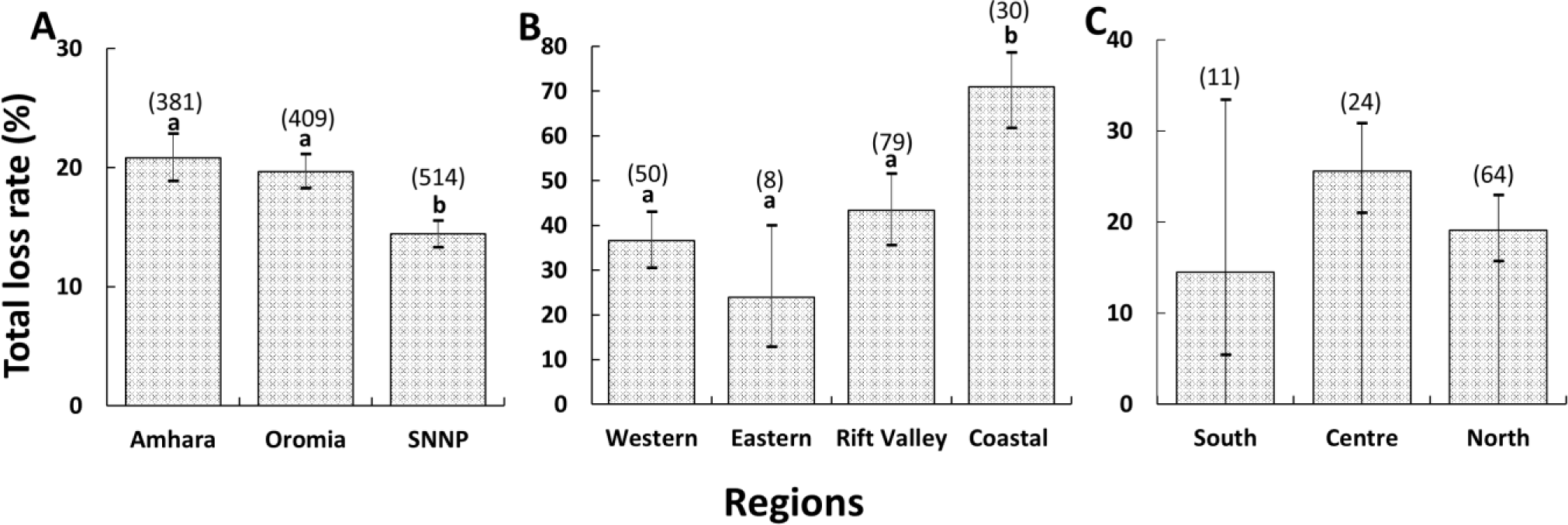
Bar graphs showing the total loss rate and 95% confidence interval in each region in Ethiopia (**A**), Kenya (**B**), and Benin (**C**). Different letters above bars indicate significant differences among the regions when compared using the quasibinomial GLM with region as an explanatory factor in van der Zee et al. [41], *p* < 0.05. The numbers in brackets above bars indicate the total number of interviewed beekeepers who provided valid data for each region.

### Detailed analysis of the risk factors associated with the losses across the countries and within Ethiopia, Kenya, and Benin

Considering the beekeeper-perceived causes of colony losses across the nine surveyed countries, issues beyond the beekeeper’s control (theft, drought, and bushfire), followed in descending order of significance, by absconding, pests/pathogens (wax moth, small and large hive beetle, ants, *Varroa* mite and *Nosema*), and pesticide poisoning (Figure 3A, Table S1). This order of risk factors varies among the surveyed countries. For instance, in Uganda, the order of risk factors was issues beyond the beekeeper’s control (mainly theft) first, followed by absconding, queen problems (queen loss during harvesting) and pesticide poisoning (Figure 3B). On the contrary, the three leading causes in Kenya were: issues beyond the beekeeper’s control (mainly theft and drought), absconding, and pests (wax moth, small and large hive beetles, ants and *Varroa* mite) (Figure 3C), but this pattern of risk factors varied within regions. In the Western region of Kenya, absconding (52%) and pests (16%) (mainly wax moth and the small hive beetle) majorly contributed to colony losses, whereas issues beyond the beekeeper’s control (32%) (mainly theft), pests (22%) (mainly wax moth) and queen problems (19%) (queen loss during harvesting) dominated in the Eastern region of the country. On the other hand, issues beyond the beekeeper’s control (38%) (mainly theft), pests (29%) (mainly wax moth) and absconding (23%) dominated in the Rift Valley region, whereas in the Coastal region, issues beyond the beekeeper’s control (60%) (mainly drought) and pests (32%) (mainly the large hive beetle and the wax moth) largely explained colony losses. Only 24 of the 87 Kenyan beekeepers who reported hive infestations by pests managed some of these biotic stressors. Whilst these Kenyan beekeepers did not treat their colonies against *Varroa* mite, wax moths, small and large hive beetles, they used an insecticide called “Sevin Dudu Dust” with an active ingredient of Carbaryl 7.5% against ants. Additionally, they also applied grease or waste engine oil to hive stands for ant prevention.

**Figure 3.**
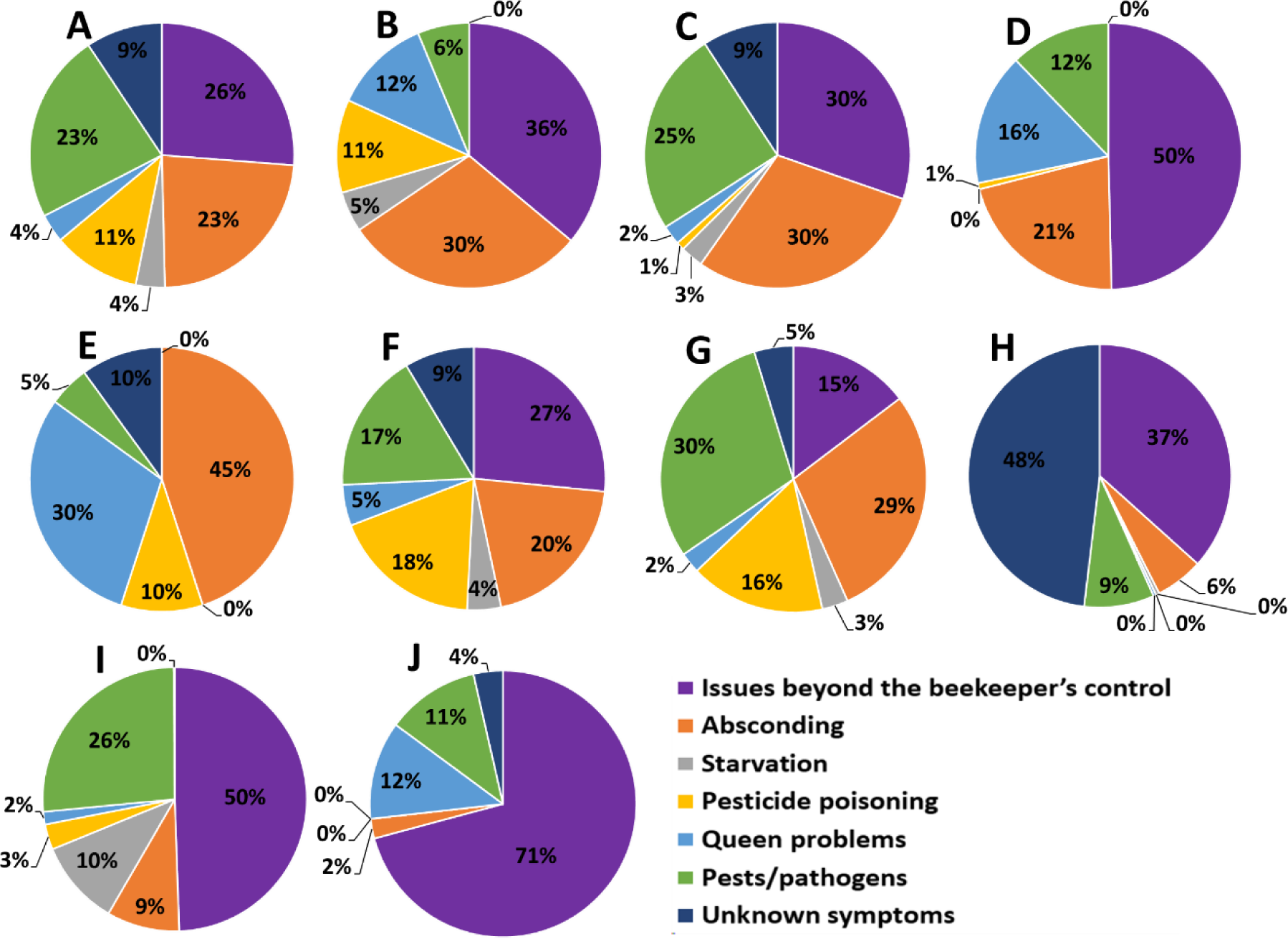
Pie charts showing the different patterns of risk factors associated with total colony losses in all the surveyed countries (**A**) and the individual countries that include Uganda (**B**), Kenya (**C**), Liberia (**D**), Nigeria (**E**), Rwanda (**F**), Ethiopia (**G**), Cameroon (**H**), Benin (**I**) and the Democratic Republic of Congo (**J**).

In Liberia, issues beyond the beekeeper’s control (mainly theft), absconding, queen problems (queen loss during harvesting) and pests (ants) explained the losses (Figure 3D). The six Liberian beekeepers who reported ant attacks treated their colonies against this pest by applying waste engine oil on hive stands. In Nigeria, absconding and queen problems (queen loss during harvesting) accounted mainly for the losses (Figure 3E), whilst in Rwanda, issues beyond the beekeeper’s control (mainly theft), absconding, pesticide poisoning, and pests (mainly ants) were responsible for the losses (Figure 3F).

In Ethiopia, the leading causes of colony losses were pests/pathogens (wax moth, small and large hive beetle, ants, *Varroa* mite and *Nosema*), absconding, pesticide poisoning, and issues beyond the beekeeper’s control (mainly theft) (Figure 3G), but the pattern of these risk factors varied across different regions. In the Amhara region, absconding was the major cause accounting for 31% of losses, followed by pesticide poisoning at 23%, and pests/pathogens at 22%. The major pests and pathogens reported by the beekeepers in this region were wax moth and the small hive beetles followed in descending order by *Varroa* mite, *Nosema* and the large hive beetle. The presence of excreta (diarrhoea) on hive components was an indication of *Nosema* infection [42]. It is worth noting that the respondents reported 31 colonies lost due to nosemosis out of the 8,973 total managed colonies from this region. This represents a total loss rate of just 0.3% out of the 20.8% loss rate reported for the Amhara region. In the Oromia region, on the other hand, pests (36%), issues beyond the beekeeper’s control (28%) (mainly theft) and absconding (25%) dominated. The major reported pests in this region were wax moths, followed by small hive beetles and *Varroa* mites. In the SNNP region, pests (39%), absconding (32%) and pesticide poisoning (21%) dominated. Beekeepers from this region reported wax moths, followed by *Varroa* and the small hive beetle, as their major pests. Whilst the Ethiopian beekeepers did not treat their colonies against *Varroa* mite and nosemosis, most of them adopted good apiary hygiene against wax moth, small and large hive beetle infestations. They also used fresh wood ash on hive stands against ants.

In Cameroon, issues beyond the beekeeper’s control (mainly theft) dominated, but most beekeepers could not explain the factors responsible for the losses (Figure 3H). Issues beyond the beekeeper’s control (mainly theft) and pests (mainly ants) dominated in Benin (Figure 3I). When looking at the pattern of risk factors associated with the colony loss rate across the regions in Benin, our results revealed that pests (56%) (small hive beetle, ants and *Varroa* mite) and issues beyond the beekeeper’s control (26%) (mainly theft) dominated in the southern region of the country. In the Central region, issues beyond the beekeeper’s control (56%) (mainly theft) and starvation (19%) majorly explained the losses, whereas issues beyond the beekeeper’s control (46%) (mainly theft) and pests (36%) (mainly ants) dominated in the Northern region. Most beekeepers in Benin who reported ant predation used permethrin insecticide and waste engine oil against ants. In the Democratic Republic of Congo, issues beyond the beekeeper’s control (mainly theft and bush fire), queen problems (queen loss during harvesting) and pests (mainly ants and termites) dominated (Figure 3J). Beekeepers in this country often used fresh wood ash, grease, or oil (e.g. vegetable oil or waste engine oil) on hive stands against ants and termites.

### Detailed analysis of the effect of management and hive types on the losses across the countries and within Ethiopia, Kenya, and Benin

Our results showed that the type of beekeeping significantly influenced the total loss rate across the nine participating countries (quasibinomial GLM: F = 37.2, df = 2, *p* < 0.001), with professional beekeepers losing fewer colonies (14.3% (95% CI: 11.0-18.3%)) than semi-professionals (22.8% (95% CI: 21.8-23.7%)) and hobbyists (39.6% (95% CI: 31.5-48.4%)). A similar result was obtained in Kenya (quasibinomial GLM: F = 10.4, df = 1, *p* < 0.01) and Benin (quasibinomial GLM: F = 4.9, df = 2, *p* < 0.01), but not in Ethiopia (GLM. quasibinomial: F = 0.7, df = 1, *p* = 0.4). Most of the respondents (94%) across the nine surveyed countries were semi-professionals, compared to 5% and 1% who were professionals and hobbyists, respectively. On average, the professional beekeepers managed 97.0 ± 31.7 (95% CI) colonies compared to 19.7 ± 1.1 (95% CI) and 19.9 ± 16.2 (95% CI) colonies managed by semi-professionals and hobbyists, respectively. Most respondents in Ethiopia (98.8%), Kenya (93%) and Benin (84%) were also semi-professionals. These semi-professional beekeepers, on average, managed 17.4 ± 0.6 (95% CI) colonies, which was approximately two-fold lower than professionals (30.1 ± 5.8 (95% CI)) in Ethiopia. In the same vein, professional beekeepers in Kenya, on average, managed 29.2 ± 20.3 (95% CI) colonies compared to 16.6 ± 3.6 (95% CI) colonies managed by semi-professionals. In Benin, the professional beekeepers on average managed 167.7 ± 165.2 (95% CI) colonies compared to 22.9 ± 4.0 (95% CI) and 14.8 ± 10.3 (95% CI) colonies managed by semi-professionals and hobbyists, respectively.

Our results further showed that the total colony loss rate for migratory beekeepers (17.9% (95% CI: 16.7-19.2%)) was significantly less than for stationary beekeepers (23.2% (95% CI: 22.0-24.5%)) across the nine participating countries (quasibinomial GLM: F = 30.7, df = 1, *p* < 0.001). These migratory beekeepers represented only 39.5% of the total respondents and on average managed 21.5 ± 3.4 (95% CI) colonies when compared to the 60.5% stationary ones, who, on average, managed 24.6 ± 2.3 (95% CI) colonies. Migratory beekeepers moved their colonies during the active beekeeping season, mainly for crop pollination and honey production. The pollinated crops were maize, sunflower, coffee, sorghum, avocado, orange, mango, pawpaw, niger seed, and macadamia. Similarly, the 45% of Ethiopian beekeepers who migrated their colonies registered significantly lower losses of 16.8% (95% CI: 15.8-17.9%), than the 55% of stationary beekeepers (19.7% (95% CI: 18.4-21%)) (quasibinomial GLM: F = 10.2, df = 1, *p* < 0.01). On average, these migratory beekeepers, who were all semi-professionals, managed 15.4 ± 0.6 (95% CI) colonies when compared to the 19.3 ± 1.0 (95% CI) colonies managed by stationary ones. In contrast, migratory and stationary beekeepers had similar losses in Kenya (quasibinomial GLM: F = 1.2, df = 1, *p* = 0.3) and Benin (quasibinomial GLM: F = 0.5, df = 1, *p* = 0.5).

In this survey, many of the respondents (79%) in the nine African countries who received training on best beekeeping practices lost significantly fewer colonies (20% (95% CI: 19.0-21.0%)) than the 21% of beekeepers who did not receive such training (26.1% (95% CI: 24.0-28.4%)) (quasibinomial GLM: F = 29.3, df = 1, *p* < 0.001). Conversely, 81%, 77% and 92% of respondents who received training in Ethiopia (quasibinomial GLM: F = 0.04, df = 1, *p* = 0.8), Kenya (quasibinomial GLM: F = 0.2, df = 1, *p* = 0.6) and Benin (quasibinomial GLM: F = 0.1, df = 1, *p* = 0.7), respectively, had similar loss rates to those who did not receive any training.

Across all the participating countries, the 76% of beekeepers who provided supplementary feeds to their colonies before the swarming season registered significantly lower losses of 18.7% (95% CI: 17.9-19.6%)) than those (24%) who did not (26.1% (95% CI: 24.0-28.4%)) (quasibinomial GLMl: F = 58.4, df = 1, *p* < 0.001). These beekeepers received training on best beekeeping practices and fed their colonies with different sugar sources (e.g. sugar syrup, Bee Fonda, mango, papaya, pineapple, and/or banana juices) and/or pollen substitutes (e.g. cassava, maize, sorghum, roasted pea, bean, soya, and/or barley flours). In the same vein, the beekeepers (82%) who provided water to their colonies registered significantly lower losses of 20.4% (95% CI: 19.5-21.4%)) than the 18% of beekeepers who did not (23.2% (95% CI: 20.9-25.7%)) (quasibinomial GLM: F = 8.1, df = 1, *p* < 0.01). Supplementary feeding and/or provision of water were most practiced by beekeepers from Ethiopia and Rwanda, whereas those from Liberia and Nigeria practiced neither of these. In fact, all the Ethiopian beekeepers interviewed provided water to their managed colonies but no supplementary feeding at the onset of the swarming season. Meanwhile, the loss rate of Kenyan beekeepers who provided supplementary feeding (quasibinomial GLMl: F = 0.4, df = 1, *p* = 0.5) or water (quasibinomial GLM: F = 0.1, df = 1, *p* = 0.8) was not significantly different from that of beekeepers who did not provide any of these. It is worth noting that only 8.4% and 32.9% of Kenyan beekeepers provided supplementary feeding and water, respectively. In Benin, the loss rate of the 7.1% of beekeepers who provided supplementary feeding (quasibinomial GLM: F = 0.0, df = 1, *p* = 1) was not significantly different from that of the 93% of beekeepers who did not. Intriguingly, the 45% of beekeepers in Benin who provided water to their colonies had a significantly higher loss rate of 25.8% (95% CI: 21.9-30%) than the 55% of beekeepers who did not (16.3% (95% CI: 13-20.4%) (quasibinomial GLM: F = 11.7, df = 1, *p* < 0.001).

The survey results also revealed that the loss rate in movable frame hives (10.6% (95% CI: 9.7-11.5%)) was significantly lower than that in local frameless hives across all the nine African countries (17.9% (95% CI: 16.5-19.3%)) (quasibinomial GLM: F = 120.8, df = 1, *p* < 0.001). More than 90% of colony migrations were carried out by beekeepers who had movable frame hives. Similarly, the loss rate in the local hives (16.1% (95% CI: 15.4-16.9%)) was approximately three-fold higher than that in the modern hives in Ethiopia (5.9% (95% CI: 5.5-6.5%)) (quasibinomial GLM: F = 481.4, df = 1, *p* < 0.001).

### Increase and decrease in the number of bee swarm catches over the past three years in Ethiopia and Kenya

As shown in Figure 6A, the number of bee swarms caught over the past three years decreased significantly by one and half-fold in Kenya (Poisson GLM: F = 18.2, df = 2, *p* < 0.001). The major decrease occurred between 2021-22 to 2022-23. In contrast, there has been a highly significant increase (∼one and half-fold) in the number of bee swarm catches over the past three years in Ethiopia (Poisson GLM: F = 99, df = 2, *p* < 0.001). When comparing the number of bee swarm catches between Ethiopia and Kenya over the past three years, we found a significant effect of year (Poisson GLM: F = 88.7, df = 2, *p* < 0.001), country (Poisson GLM: F = 142.9, df = 1, *p* < 0.001) and the interaction between the year and the country (Poisson GLM: F = 28.5, df = 2, *p* < 0.001). In fact, bee swarm catches were not the same among the years and were one and half-fold higher in Kenya than in Ethiopia. The number of bee swarm catches differed within the individual surveyed regions of Ethiopia (Figure 2B). While a significant decrease (∼one- and one and half-fold) in the number of bee swarm catches was found in the Amhara region from the year 2020/2021 to the year 2021/2022 (Poisson GLM: F = 80.7, df = 2, *p* < 0.001), the contrary was observed in the Oromia region. In this region, the number of bee swarms caught increased significantly by approximately one and half-fold more in the year 2022/2023 than in the years 2021/2022 and 2020/2021 (Poisson GLM: F = 58.7, df = 2, *p* < 0.001). Similarly, the number of bee swarms caught differed significantly between the years in the SNNP region (Poisson GLM: F = 76.7, df = 2, *p* < 0.001), increasing by approximately one- or one and half-fold more in the year 2022/2023 than the years 2021/2022 or 2020/2021, respectively. Similarly, when comparing the number of bee swarm catches among regions in Ethiopia over the past three years, we found a significant effect of year (Poisson GLM: F = 99.0, df = 2, *p* < 0.001), region (Poisson GLM: F = 421.3, df = 1, *p* < 0.001) and the interaction between the year and the region (Poisson GLM: F = 58.6, df = 2, *p* < 0.001). In fact, bee swarm catches were not the same among the years and were higher in Amhara than Oromia and SNNP regions over the past two years.

**Figure 6.**
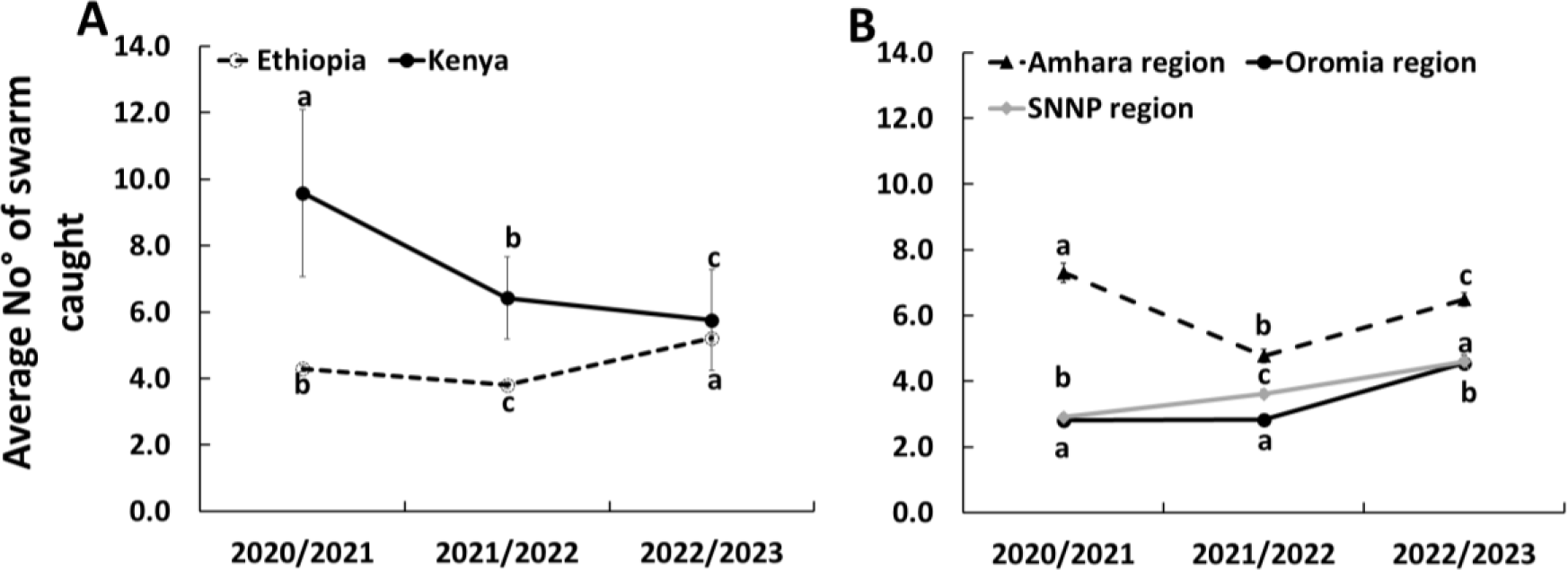
Line graph showing the average increase or decrease in the number of bee swarms caught over the past two years (Mean ± SE) in Ethiopia and Kenya (**A**) and in Amhara, Oromia and SNNP regions of Ethiopia (**B**). Different letters above bars indicate significant differences among the years within the region when compared using the Poisson GLM with log link.

## Discussion

The sustainability of apiculture and pollination services depends on the ability of the beekeepers to maintain at least the same number of productive colonies over time. This survey revealed that the majority of interviewed beekeepers (95.3%) rely on capturing bee swarms to compensate for their colony losses and even expand their apiary size, as is generally known in Africa [29]. Apparently, the sustainability of this practice is at risk in Kenya and in the Amhara region of Ethiopia due to a significant decrease in bee swarm availability over the past three years. An opposite trend over the same length of time was observed in the Oromia and SNNP regions of Ethiopia. This disparity might be attributed to differential impacts of environmental factors such as land use change, pesticide use and/or climate change on the availability of wild and managed honey bee populations, which are potential sources of bee swarms during the swarming season. These environmental stressors have been recently identified as major contributors to pollinator decline in Africa [43]. This declining trend in swarm availability in Kenya was initially reported in 2010 by Muli et al. [44]. Similar observations were made between 2005-2006 in the Amhara region of Ethiopia, mainly due to the expansion of farmland areas and pesticide use [45]. Our findings further support the significance of pesticide poisoning in colony losses in Ethiopia (Figure 3G), particularly in the Amhara and SNNP regions. Reports indicate that pesticide usage among farmers in the Amhara and SNNP regions is common due to the heavy production of cereal, vegetable, and fruit crops [45–47]. In fact, the use of pesticides known to harm beneficial insects, including honey bees, has been documented not only in Ethiopia [45,48–50], but also in Kenya [51–53] and other parts of Africa [55]. These pesticides include organochlorines (e.g. endosulfan and dichlorodiphenyltrichloroethane (DDT)) and neonicotinoids (e.g. imidacloprid, thiamethoxam, acetamiprid). Given these findings, we recommend the implementation of long-term spatio-temporal studies in Kenya, Ethiopia, and other African countries to better understand the dynamics of swarm availability and to pinpoint the underlying causes. Such research is essential for informed decision-making and effective strategic planning in the field of apiculture.

Regarding the losses, our study revealed that the overall loss rate of honey bee colonies varied significantly among the nine participating African countries during the active beekeeping season, ranging from 9.7-45.3%. This season corresponds to the spring and summer periods in the Southern hemisphere [56]. Loss rates were highest at 42-45% in Uganda and Kenya, whilst the lowest rates at 9.7-14.6% were in Liberia, Nigeria, and Rwanda. The overall loss rates also differed considerably among regions within Ethiopia and Kenya and were in the range of 14.4-20.8% and 23.9-71%, respectively. Further, regions of higher and lower risk of losses were observed within each country when calculating the relative risk of total colony losses at regional level (Figure 7). The range of total colony losses recorded herein was similar to those reported for South African honey bees during the active beekeeping season (29.6-46.2%) [20]. It is important to note that respondents expressed concerns about their levels of losses, as these losses pose a threat to the viability of their businesses. The considerable variations in the above loss rates among African countries could be attributed to honey bee genetic characteristics, pests, environmental and sociological factors, as well as colony management practices.

**Figure 7.**
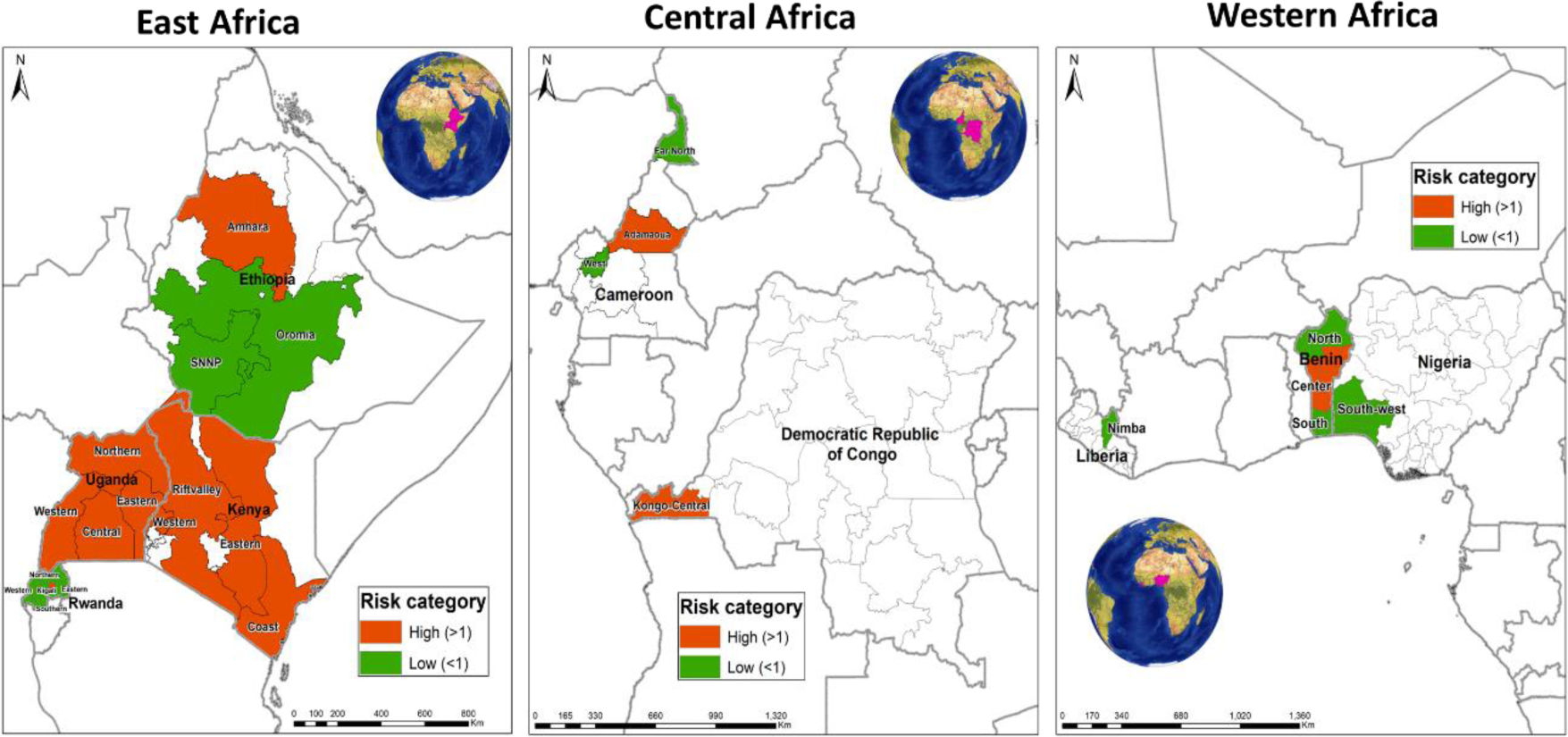
Map with colour coding showing the relative risk of loss at the regional level for each country in Sub-Saharan Africa. The relative risk of loss for each region was calculated as the loss rate for that region divided by the loss rate of all regions as, in Gray et al. [14]. Red and green colours indicate regions with a relative risk of loss higher and lower than one, respectively. All regions considered for this analysis had at least three beekeepers.

Risk factors affecting apiculture can be divided into external and internal. Among the external factors, issues beyond the beekeeper’s control and pesticides may have played a major role. Other critical internal factors absconding, pests, colony management practices and sociological factors. Among the issues beyond the beekeeper’s control, three external risk factors reported by the beekeepers were the most pronounced in order of significance: theft, drought, and bushfire. Their impact was most pronounced in the Democratic Republic of Congo (71%) (Figure 2J), with no impact in Nigeria (Figure 2E). To address the significant issue of thefts, initiatives such as the development of owner-specific branded hives/frames, provision of hive insurance, development of state laws imposing heavy penalties for stealing beehives, and/or investment in anti-theft technologies (e.g. GPS trackers and/or surveillance cameras) may help in the future, as was suggested for similar cases in South Africa by Masehela [57]. The second significant issue beyond the beekeeper’s control was a prolonged drought in 2022, and this was particularly pronounced in the semi-arid regions of Kenya (i.e. Eastern, Rift Valley and Coastal regions) and the Central and Northern regions of Benin. This natural disaster event may limit forage and water availability for honey bees [58] as well as supporting bush fires [59]. Bush fires, caused by slash-and-burn agriculture to prepare for the next planting season, were the third important issue beyond the beekeeper’s control. The practice was particularly significant in the Democratic Republic of Congo. To date, farmers in developing countries in Asia and Africa still practice slash-and-burn agriculture [60]. While climatic events like drought are unpredictable [60,61], strategies to mitigate the impact of widespread bushfires should be developed and enforced, especially under drought conditions, as was suggested for similar issues in Uganda [63]. These efforts could include promoting sustainable agricultural practices that reduce the reliance on slash-and-burn methods, enhancing early warning systems for drought and fire, and improving water resource management to support apiculture during dry periods [63].

As highlighted above, absconding significantly contributed to the total losses reported in this study, exhibiting the highest impact in Nigeria (45%) and the lowest in the Democratic Republic of Congo (2%). Absconding, a non-reproductive form of swarming where all adult colony members including the queen vacate the hive, was also identified as a cause of colony losses during the active beekeeping season in South Africa [20]. It is worth noting that absconding is considered as a pronounced behaviour of the African honey bee subspecies in response to stress caused by various disturbances [23,63,64], but subspecies’ tendency to abscond varies [29]. Besides the honey bee genetics [66], several factors can induce absconding, including pests and predation, human manipulation and limited resources [63,66]. For instance, some pests contributing to colony losses in this study were previously reported to elicit colony absconding. These include the wax moth [67,68], the small hive beetle [70], the large hive beetle [71] and ants [28,71]. Of note, attack of a hive by the large hive beetles was widespread in the Coastal region of Kenya, where an exceptionally high infestation rate of the large hive beetle, with over 400 individuals per hive, has recently been documented [29,70]. On the other hand, ant predation was predominantly observed in Rwanda, Cameroon, the Democratic Republic of Congo, the Amhara and Oromia regions of Ethiopia, and the Southern and Northern regions of Benin. Additionally, external environmental factors such as drought conditions reported above and starvation felt mainly by beekeepers in the Central region of Benin may also trigger the relocation of honey bee colonies to areas with more favourable microclimatic conditions and/or forage resources, as seen in some ant species [73]. Furthermore, even routine human activities such as colony examinations can precipitate absconding [63,73], highlighting the complex interplay of biotic and abiotic stresses impacting bee colony stability. It should be noted that while absconding colonies are a personal loss to the beekeeper, they are not lost to the ecosystem and pollination activity.

Regarding the role of colony pests and pathogens in colony loss, as reported in our survey, the respondents mainly reported the pests visible to their naked eyes. Among the pathogens, only Nosema infection was reported, but it is important to indicate that this infection was detected indirectly by fecal spots and was not confirmed by any microscopic or molecular analysis. However, the occurrence of Nosema was not widespread but rather confined to managed colonies in the Amhara region of Ethiopia. Previous studies have documented *Nosema* infection caused by *N. apis* in Ethiopia [75–77], but the extent to which it affects the health of honey bees, the apicultural industry, and its mode of management, still remain largely unclear in Ethiopia. In general, participants of the survey have limited knowledge on the identification and management of pathogens that can compromise honey bee health and productivity. This knowledge gap underscores the urgent need to enhance local capacities in pathogen diagnosis to safeguard the health of African honey bees. Strengthening these capabilities would involve training beekeepers and local researchers in advanced diagnostic techniques, increasing awareness of bee health issues, and encouraging the adoption of best management practices to mitigate pathogen impact. Such initiatives are crucial for the sustainable development of apiculture in Africa.

In our survey, all the pests and predators reported as risk factors are generally considered benign to African honey bee colonies [15]. For instance, *Varroa* mite was reported not significant in a few of the surveyed countries such as Kenya [35–37], Ethiopia [39] and Uganda [78]. This is also true in South Africa [15] and in Tunisia [79]. However, the impact of the mite on managed honey bee colonies in other surveyed countries (e.g. Rwanda, Cameroon, Democratic Republic of Congo, Benin, Nigeria, and Liberia) remains unknown. In the same vein, the adoption of good apiary hygiene practices generally helps to minimize damages and colony losses attributed to pests like the wax moth, ants, small and large hive beetles [15,68,80]. Nevertheless, the beekeepers in most surveyed countries mentioned *Varroa*, wax moths, ants, small and large hive beetles as threats to their business. Since these pests are visually conspicuous, this could be a misattribution, blaming visible symptoms rather than underlying causes. In particular, small hive beetles and wax moths, which scavenge on abandoned hive resources, are often mistakenly identified as the primary cause of colony losses rather than a symptom of deeper management issues. The prevalence of such perceptions among these respondents may be related to poor apiary management practices, as over 90% of them are part-time (semi-professional) beekeepers who lose more colonies compared to their full-time (professional) counterparts. This observation suggests a potential correlation between beekeeping engagement level and pest impact, warranting further investigation to clarify these dynamics and improve colony management strategies. Although the survey did not explicitly capture the impact of hive types on colony losses due to these pests, our results indicated that losses were significantly higher in locally made hives than the modern ones. This finding underscores the need to explore further how indigenous hive types and management practices influence the rate of losses due to pests, pathogens, and other factors.

The survey revealed that most respondents who did not engage in migratory beekeeping experienced considerably lower losses overall than those who did. This pattern was consistent in Ethiopia, but not in Kenya and Benin, where losses were similar between migratory and non-migratory beekeepers. These varying results could be explained by differences in the quality and/or quantity of forage resources available to migratory and stationary colonies within their respective landscapes. In the USA [81], Austria (in 2019) [82] and across Europe (in 2018 and 2020) [4,14], a lower loss rate in migratory than non-migratory beekeeping was observed, whereas an opposite trend was observed in South Africa [20] and Europe (in 2019) [5].

The survey results also showed a positive association between colony loss rate and supplementary feeding and/or provision of water within an African context for the first time. Over 70% of the interviewed beekeepers fed their colonies at the onset of swarming to aid colony establishment and/or to stimulate comb construction and population build-up. This management practice apparently contributed to lower colony losses across the surveyed countries, except in Kenya and Benin, where no favourable impact was reported. This finding is not surprising as supplement types and quantities vary between the countries, and may affect colony performance and viability differently, as previously demonstrated [83–86]. For example, in Kenya and Benin, 8.4% and 7.1% of respondents, respectively, used remains of honey in beeswax to feed their colonies. In contrast, in Ethiopia and Rwanda where feeding was most prevalent, a mixture of carbohydrate sources (e.g. sugar syrup, Bee Fonda, mango, papaya, pineapple, and/or banana juices) and pollen substitutes (e.g. cassava, maize, sorghum, roasted pea, bean, soya, and/or barley flours) were often used. Since the type of protein affects the physiology of the honey bees [87,88], the observed variability could result from the different protein sources. The contribution of these different feeds to colony performance and health needs to be studied in the future. Similarly, the impact of water provision at the onset of swarming varied between the countries, but it did not affect the colony loss rate in Kenya, whereas beekeepers in Benin who provided water to their colonies had higher losses than those who did not. The latter finding in Benin could be a result of water contamination in a way that spreads harmful agents within the colonies [89], but this is surprising and requires further investigation.

Sociological factors, particularly education in beekeeping, were also related to the colony loss rate in countries such as Liberia, Nigeria, Rwanda, and Ethiopia, which recorded considerably lower losses (9.7 - 18.6%) and had more beekeepers who received training on best beekeeping practices (45 - 80.6%) than Cameroon, which recorded a 27.9% loss rate while only 26% of respondents received training in beekeeping (Figure 1). These findings align with previous reports indicating a lack of training in bee farming in Cameroon [90]. Overall, these results underscore the need to educate the beekeepers on identifying and managing potential risk factors that affect managed colony health and productivity in this country.

### Conclusions

This pioneering study provides an inaugural examination of managed honey bee colony loss rates and identifies several key risk factors within nine Sub-Saharan African countries during the active beekeeping season. Despite the sample size covering only one percent of all estimated beekeepers in each participating country, the implications of this research are significant, offering new insights into the challenges that the apicultural sector faces in this continent. The findings underscore critical risk factors that potentially threaten the sustainability of beekeeping, including pest and disease management, environmental stressors, and beekeeping practices. These challenges highlight the vulnerability of the apicultural sector in Africa, which is vital for pollination services that support agriculture and biodiversity. Given the study’s findings, there is a pressing need for ongoing, regular, long-term monitoring of honey bee colony losses in Africa and the variables influencing them. Implementing systematic and regular monitoring initiatives across the continent will help understand broader patterns and causes of bee colony declines and facilitate the development of targeted interventions to mitigate these losses. Such efforts will be crucial in fostering a resilient and sustainable apicultural sector in Africa, ensuring the health of honey bee populations and by extension, securing the ecological benefits they provide, as well as helping to secure the livelihoods of beekeepers. As the global importance of pollinators continues to gain recognition, enhancing the stability of honey bee colonies in Africa becomes a regional priority and a global one.

## Author Contributions

### Conceptualization of the research

Beatrice T. Nganso, Victoria Soroker, Alison Gray, Robert Brodschneider, Sevgan Subramanian.

### Investigation

Beatrice T. Nganso, Workneh Ayalew, Abebe J. Wubie, Freweini Assefa, Lulseged Belayhun, Nelly N. Ndungu, Daniel Toroitich, Z. Ngalo Otieno-Ayayo, Mbatha B. Wambua, Yudah O. Oyieyo, Ntirenganya Elie, Rachidatou Sikirou, Souradji B. Idrissou, Willy Mwiza, S. Turner, Bridget O. Bobadoye, Sidonie T. Fameni, Sayemie Gaboe, Mawufe K. Agbodzavu, Patrick Mafwila, Geraud C. Tasse Taboue.

### Resources

Beatrice T. Nganso, W. A., Nelly N. Ndungu, Z. Ngalo Otieno-Ayayo, Mawufe K. Agbodzavu, Ntirenganya Elie, Souradji B. Idrissou, Rachidatou Sikirou, S. Turner, Bridget O. Bobadoye, Sidonie T. Fameni, Sayemie Gaboe.

### Data curation

Beatrice T. Nganso

### Data analysis

Beatrice T. Nganso, Kimathi Emily

### Writing of original draft

Beatrice T. Nganso.

### Supervision

Beatrice T. Nganso, Victoria Soroker, Alison Gray, Robert Brodschneider.

### Funding acquisition

Beatrice T. Nganso, Workneh Ayalew, Nelly N. Ndungu, Sevgan Subramanian, Rachidatou Sikirou.

### Editing the manuscript

All authors have read and agreed to the published version of the manuscript.

## Funding

Data collection on colony losses in Kenya was funded by the European Union project through Kenya Agricultural and Livestock Research Organization (KALRO), Grant/Award Number: KALRO/CS APP/LOA No. 3/2019; whereas in Ethiopia, it was funded by the Mastercard Foundation through the More Young Entrepreneurs in Silk and Honey (MOYESH) Programme. In Cameroon, data collection was funded by JRS Biodiversity Foundation, Grant Number: 70054, whereas in Rwanda and Benin, it was funded by the COLOSS Panuwan Chantawannakul Award 2023 to Dr. Beatrice T. Nganso.

## Data Availability Statement

The data presented in this study are available on request from the corresponding author.

## Acknowledgements

The authors acknowledge all beekeepers for generously providing responses needed for this survey. The authors are also grateful to the COLOSS research association (prevention of honey bee colony losses) which facilitated increased data collection in Rwanda and Benin through one of its funding agencies, the Ricola Foundation. The authors further acknowledge the financial support for this research by the following organizations and agencies: the Swedish International Development Cooperation Agency (Sida); the Swiss Agency for Development and Cooperation (SDC); the Australian Centre for International Agricultural Research (ACIAR); the Norwegian Agency for Development Cooperation (Norad); the German Federal Ministry for Economic Cooperation and Development (BMZ); and the Government of the Republic of Kenya. The views expressed herein do not necessarily reflect the official opinion of the donors.

## Conflicts of Interest

The authors declare that they have no competing interests.

---

**Table S1.** Survey results for 2022/2023, showing the total number of respondents with valid loss data per country, corresponding number of managed honey bee colonies during the active beekeeping season, rates of loss due to natural disasters, absconding, starvation, poor harvesting techniques, pesticide poisoning, predators, queen problems, pests/diseases/pathogens, other factors and unknown problems, and overall loss rate (each with 95% confidence intervals (CIs))

| Countries | No of respondents | No of managed colonies | % rate of loss of colonies due to issues beyond the beekeeper's control (95% CI) | % rate of loss of colonies due to absconding (95% CI) | % rate of loss of colonies due to starvation (95% CI) | % rate of loss of colonies due to pesticide poisoning (95% CI) | % rate of loss of colonies due to queen problems (95% CI) | % rate of loss of colonies due to pests/pathogens (95% CI) | % rate of loss of colonies due to unknown problems (95% CI) | % total loss rate (95% CI) |
| --- | --- | --- | --- | --- | --- | --- | --- | --- | --- | --- |
| **East Africa** |  |  |  |  |  |  |  |  |  |  |
| Ethiopia | 1304 | 22,838 | 2.7 (2.4-3.0) | 5.3 (4.9-5.8) | 0.6 (0.5-0.7) | 3.1 (2.7-3.4) | 0.5 (0.4-0.6) | 5.5 (5.1-6.0) | 0.9 (0.7-1.1) | 18.6 (17.7-19.5) <sup>b</sup> |
| Kenya | 167 | 2,875 | 12.9 (10.2-16.1) | 12.5 (10.5-14.7) | 1.2 (0.8-1.8) | 0.4 (0.2-0.7) | 1.0 (0.6-1.6) | 10.6 (8.7-12.7) | 3.9 (2.7-5.7) | 42.4 (37.8-47.2) <sup>d</sup> |
| Rwanda | 58 | 5,573 | 3.9 (2.1-7.1) | 2.9 (1.7-5.0) | 0.6 (0.3-1.5) | 2.7 (1.3-5.3) | 0.7 (0.3-1.7) | 2.5 (1.0-6.0) | 1.3 (0.5-3.3) | 14.6 (10.4-20.1) <sup>a</sup> |
| Uganda | 25 | 1,341 | 16.3 (12.4-21.2) | 13.4 (9.8-17.9) | 2.2 (1.0-4.7) | 5.2 (2.6-9.8) | 5.4 (2.6-10.6) | 2.8 (1.1-7.2) | 0.0 (0.0-0.0) | 45.3 (36.8-54.1) <sup>d</sup> |
| **West Africa** |  |  |  |  |  |  |  |  |  |  |
| Benin | 99 | 3,665 | 10.5 (8.4-12.9) | 1.9 (1.1-3.2) | 2.2 (1.6-3.1) | 0.6 (0.3-1.3) | 0.3 (0.1-0.9) | 5.6 (3.9-7.8) | 0.0 (0.0-1.2) | 21.1 (19.2-23.2) <sup>bc</sup> |
| Liberia | 18 | 1,354 | 4.8 (2.4-9.5) | 2.1 (0.9-4.5) | 0 | 0.1 (0.0-3.8) | 1.5 (0.6-3.7) | 1.2 (0.3-4.6) | 0.0 (0.0-0.0) | 9.7 (4.8-18.6) <sup>a</sup> |
| Nigeria | 20 | 169 | 0 | 5.3 (2.9-9.5) | 0 | 1.2 (0.2-7.7) | 3.5 (1.3-9.6) | 0.6 (0.1-2.7) | 1.2 (0.3-4.3) | 11.8 (6.8-19.8) <sup>abc</sup> |
| **Central Africa** |  |  |  |  |  |  |  |  |  |  |
| Democratic Republic of Congo | 49 | 655 | 18.2 (13.7-23.7) | 0.6 (0.1-4.2) | 0 | 0 | 3.0 (1.3-7.2) | 2.9 (1.3-6.4) | 0.9 (0.4-2.0) | 25.6 (20.8-31.2) <sup>bc</sup> |
| Cameroon | 46 | 3,291 | 10.2 (8.5-12.2) | 1.6 (0.8-3.2) | 0.1 (0.0-1.2) | 0 | 0.1 (0.0-0.3) | 2.4 (1.4-4.0) | 13.4 (10.0-17.7) | 27.9 (23.1-33.2) <sup>c</sup> |
| **Total** | 1,786 | 41,761 | 5.6 (5.1-6.1) | 5.0 (4.6-5.4) | 0.7 (0.6-0.9) | 2.3 (2.0-2.6) | 0.7 (0.6-0.9) | 5.0 (4.5-5.4) | 2.0 (1.7-2.3) | 21.3 (20.4-22.2) |

\*Different superscript letters (<sup>a</sup>, <sup>b</sup>, <sup>c</sup>, <sup>d</sup> and <sup>bc</sup>) above letters indicate significant difference in total loss rate among the countries by fitting the quasibinomial GLM with country as an explanatory factor in Van Der Zee et al. (2013), $p < 0.05$ .

## Notes

### Competing Interest Statement

The authors have declared no competing interest.

## References

[1] van der Zee R, Pisa L, Andonov S, Brodschneider R, Charrière JD, Chlebo R, et al. Managed honey bee colony losses in Canada, China, Europe, Israel and Turkey, for the winters of 2008–9 and 2009–10. J. Apic. Res. 2012; 51(1):100–114. 10.3896/IBRA.1.51.1.12

[2] VanEngelsdorp D, Hayes Jr J, Underwood RM, Pettis J. A survey of honey bee colony losses in the US, fall 2007 to spring 2008. PloS one. 2008; 3(12): e4071. 10.1371/journal.pone.0004071

[3] Ferland J, Kempers M, Kennedy K, Kozak P, Lafrenière R, Kozak P, et al. Canadian Association of Professional Apiculturists statement on honey bee wintering losses in Canada. Canadian Association of Professional Apiculturists (CAPA). 2022; pp. 1–24.

[4] Gray A, Brodschneider R, Adjlane N, Ballis A, Brusbardis V, Charrière JD, et al. Loss rates of honey bee colonies during winter 2017/18 in 36 countries participating in the COLOSS survey, including effects of forage sources. J. Apic. Res. 2019; 58(4):479–85. 10.1080/00218839.2019.1615661

[5] Gray A, Adjlane N, Arab A, Ballis A, Brusbardis V, Charrière JD, et al. Honey bee colony winter loss rates for 35 countries participating in the COLOSS survey for winter 2018–2019, and the effects of a new queen on the risk of colony winter loss. J. Apic. Res. 2020;59(5):744–751. 10.1080/00218839.2020.1797272.

[6] Osterman J, Aizen MA, Biesmeijer JC, Bosch J, Howlett BG, Inouye DW, et al. Global trends in the number and diversity of managed pollinator species. Agric. Ecosyst. Environ. 2021;322:107653–107666. 10.1016/j.agee.2021.107653

[7] Bruckner S, Wilson M, Aurell D, Rennich K, Vanengelsdorp D, Steinhauer N, et al. A national survey of managed honey bee colony losses in the USA: results from the Bee Informed Partnership for 2017–18, 2018–19, and 2019–20. J. Apic. Res. 2023;62(3):429–443. 10.1080/00218839.2022.2158586

[8] Ollerton J, Winfree R, Tarrant S. How many flowering plants are pollinated by animals?. Oikos. 2011;120(3):321–326. 10.1111/j.1600-0706.2010.18644.x

[9] Potts SG, Imperatriz-Fonseca V, Ngo HT, Aizen MA, Biesmeijer JC, Breeze TD, et al. Safeguarding pollinators and their values to human well-being. Nature. 2016; 540:220–229. 10.1038/nature20588

[10] Popovska Stojanov D, Dimitrov L, Danihlík J, Uzunov A, Golubovski M, Andonov S, et al. Direct economic impact assessment of winter honeybee colony losses in three European countries. Agriculture. 2021;11(5):398–409. 10.3390/ agriculture11050398

[11] Vanengelsdorp D, Caron D, Hayes J, Underwood R, Henson M, Rennich K, et al.. A national survey of managed honey bee 2010–11 winter colony losses in the USA: results from the Bee Informed Partnership. J. Apic. Res. 2012;51(1):115–24. 10.3896/IBRA.1.51.1.14

[12] Brodschneider R, Gray A. How COLOSS monitoring and research on lost honey bee colonies can support colony survival. Bee World. 2022;99(1):8–10. 10.1080/0005772x.2021.1993611

[13] Lamas ZS, Chen Y, Evans JD. Case Report: Emerging losses of managed honey bee colonies. Biology. 2024;13(2):117–128. 10.3390/biology13020117

[14] Gray A, Adjlane N, Arab A, Ballis A, Brusbardis V, Bugeja Douglas A, et al. Honey bee colony loss rates in 37 countries using the COLOSS survey for winter 2019–2020: the combined effects of operation size, migration and queen replacement. J. Apic. Res. 2023;62(2):204–210. 10.1080/00218839.2022.2113329

[15] Pirk CW, Strauss U, Yusuf AA, Démares F, Human H. Honeybee health in Africa—a review. Apidologie. 2016; 47:276–300. 10.1007/s13592-015-0406-6

[16] Hristov P, Shumkova R, Palova N, Neov B. Factors associated with honey bee colony losses: A mini-review. Vet. Sci. 2020;7(4):166–183. 10.3390/vetsci7040166

[17] Potts SG, Biesmeijer JC, Kremen C, Neumann P, Schweiger O, Kunin WE. Global pollinator declines: trends, impacts and drivers. Trends Ecol. Evol. 2010;25(6):345–353. 10.1016/j.tree.2010.01.007

[18] Castilhos D, Bergamo GC, Gramacho KP, Gonçalves LS. Bee colony losses in Brazil: a 5-year online survey. Apidologie. 2019; 50:263–272. 10.1007/s13592-019-00642-7

[19] Requier F, Leyton MS, Morales CL, Garibaldi LA, Giacobino A, Porrini MP, et al. First large-scale study reveals important losses of honey bee and stingless bee colonies in Latin America. Research Square [Preprint]. 2023. 10.21203/rs.3.rs-3378800/v1

[20] Pirk CW, Human H, Crewe RM, VanEngelsdorp D. A survey of managed honey bee colony losses in the Republic of South Africa–2009 to 2011. J. Apic. Res. 2014;53(1):35–42. 10.3896/IBRA.1.53.1.03

[21] Moustafa AM, Mahbob MA, Abdel-Rahman MF, Mabrouk MS. Estimate the losses of honey bee colonies and their potential causes within the beekeepers at new valley governorate during two years survey by using questionnaire method. J. Plant Prot. Pathol. 2014; 5(3): 327–340. 10.21608/jppp.2014.87922.

[22] Underwood RM, Traver BE, López-Uribe MM. Beekeeping management practices are associated with operation size and beekeepers’ philosophy towards in-hive chemicals. Insects. 2019; 10(1): 10–23. 10.3390/insects10010010.

[23] Hepburn HR, Radloff SE. Honeybees of Africa. Springer Verlag, Berlin, Heidelberg, New York. 1988.

[24] Ilyasov RA, Lee ML, Takahashi JI, Kwon HW, Nikolenko AG. A revision of subspecies structure of western honey bee *Apis mellifera*. Saudi J. Biol. Sci. 2020; 27 (12):3615– 3621. 10.1016/j.sjbs.2020.08.001.

[25] Dietemann V, Pirk CWW, Crewe R. Is there a need for conservation of honey bees in Africa? Apidologie. 2009; 40(3):285–295. 10.1051/apido/2009013.

[26] Okwee-Acai J, Anyanzo TA, Aroba J, Vuchiri JK, Onzivua T, Okullo P. Effects of apiary management on colonisation and colony performance of African honey bee ( *Apis mellifera*) in the North-Western Agro-ecological zone of Uganda. Livest. Res. Rural Dev. 2010; 22(5): 1–9.

[27] Kuboja N, Kilima F, Isinika A. Absconding of honey bee colonies from beehives: underlying factors and its financial implications for beekeepers in Tanzania. Int. J. Agric. Sci. Res. Technol. Ext. Educ. Syst. 2020; 10(4): 185–193.

[28] Nurie YA. Factors affecting bee colony absconding and prevention mechanism in Ethiopia. Int. J. Agric. Innov. Res. 2020; 9 (2):111–119.

[29] Nganso BT, Soroker V, Osabutey AF, Pirk CW, Johansson T, Elie N, et al. Best practices for colony management: a neglected aspect for improving honey bee colony health and productivity in Africa. J. Apic. Res. 2024; 1–18. 10.1080/00218839.2024.2308418.

[30] Hailu TG, Wakjira K, Gray A. Honey bee colony population annual dynamics in northern Ethiopia’s semi-arid region, Tigray. J. Apic. Res. 2024;1–10. 10.1080/00218839.2024.2308418.

[31] Gratzer K, Wakjira K, Fiedler S, Brodschneider R. Challenges and perspectives for beekeeping in Ethiopia. A review. Agron. Sustain. Dev. 2021; 41(4): 1–5. 10.1007/s13593-021-00702-2.

[32] Ruttner F. Biogeography and taxonomy of honey bees. Springer Science & Business Media 1988.

[33] Ambaw M, Teklehaimannot T, Workye M. The prevalence of wax moth and associated risk factors in selected districts of Arsi Zone. J. Entomol. Zool. Stud. 2020;8(1):200–205.

[34] Strauss U, Dietemann V, Human H, Crewe RM, Pirk CW. Resistance rather than tolerance explains survival of savannah honey bees (*Apis mellifera scutellata*) to infestation by the parasitic mite *Varroa destructor*. Parasitology. 2016;143(3):374–87 10.1017/S0031182015001754.

[35] de Souza FS, Allsopp MH, Martin SJ. Deformed wing virus prevalence and load in honey bees in South Africa. Arch. Virol. 2021; 166(1): 237–241, 2021, 10.1007/s00705-020-04863-5.

[36] Nganso BT, Fombong AT, Yusuf AA, Pirk CW, Stuhl C, Torto B. Hygienic and grooming behaviors in African and European honey bees - New damage categories in *Varroa destructor*. PLoS One. 2017;12(6):e0179329: 10.1371/journal.pone.0179329.

[37] Nganso BT, Fombong AT, Yusuf AA, Pirk CW, Stuhl C, Torto B. Low fertility, fecundity and numbers of mated female offspring explain the lower reproductive success of the parasitic mite *Varroa destructor* in African honey bees. Parasitology. 2018; 145(12):1633–1639. 10.1017/S0031182018000616

[38] Cheruiyot SK, Lattorff HMG, Kahuthia-Gathu R, Mbugi JP, Muli E. Varroa-specific hygienic behavior of Apis mellifera scutellata in Kenya. Apidologie. 2018; 49(4):439–449. 10.1007/s13592-018-0570-6.

[39] Haftom Gebremedhn HG, Bezabeh Amssalu BA, Smet LD, Graaf DD. Factors restraining the population growth of *Varroa destructor* in Ethiopian honey bees (*Apis mellifera simensis*). PLoS One. 2019; 14(9): e0223236. 10.1371/journal.pone. 0223236.

[40] R Development Core Team. R: A language and environment for statistical computing. R Foundation for Statistical Computing, Vienna, Austria. 2023.

[41] van der Zee R, Gray A, Holzmann C, Pisa L, Brodschneider R, Chlebo R, et al. Standard survey methods for estimating colony losses and explanatory risk factors in *Apis mellifera*. J. Apic. Res. 2013;52(4):1–36. 10.3896/IBRA.1.52.4.18.

[42] Fries I, Chauzat MP, Chen YP, Doublet V, Genersch E, Gisder S, et al. Standard methods for Nosema research. J. Apic. Res.2013;52(1):1–28. 10.3896/IBRA.1.52.1.14.

[43] Dicks LV, Breeze TD, Ngo HT, Senapathi D, An J, Aizen MA, et al. Risks associated with pollinator decline. Nat. Ecol. Evol. 2021; 5(10):1453–1461. 10.1038/s41559-021-01534-9.

[44] Muli E, Patch H, Frazier M, Frazier J, Torto B, Baumgarten T, et al. Evaluation of the distribution and impacts of parasites, pathogens, and pesticides on honey bee ( *apis mellifera*) populations in East Africa. PLoS One. 2014; 9(4):e94459. 10.1371/journal.pone.0094459.

[45] Ejigu K, Gebey T, Preston TR. Constraints and prospects for apiculture research and development in Amhara region, Ethiopia. Livest. Res. Rural Dev. 2009;21(10): 172.

[46] Asmare BA, Freyer B, Bingen J. Pesticide use practices among female headed households in the Amhara Region, Ethiopia. Sustain. 2022; 14(22):1–26. 10.3390/su142215215.

[47] Damte T. Trends in pesticide use by smallholder farmers on ‘meher’ season field and horticultural crops in Ethiopia. J. Agric. Sci. 2022;32(2):133–156.

[48] Mullié WC, Prakash A, Müller A, Lazutkaite E. Insecticide use against desert locust in the Horn of Africa 2019–2021 reveals a pressing need for change. Agronomy. 2023;13(3):819–842. 10.3390/agronomy13030819.

[49] Nigatu AW, Bråtveit M, Moen BE. Self-reported acute pesticide intoxications in Ethiopia. BMC Public Health. 2016;16(1):1–8. 10.3390/su142215215.

[50] Mekonnen B, Siraj J, Negash S. Determination of pesticide residues in food premises using quechers method in bench-sheko zone, Southwest Ethiopia. Biomed. Res. Int. 2021; 19:1–3. 10.1155/2021/6612096.

[51] Negatu B, Dugassa S, Mekonnen Y. Environmental and health risks of pesticide use in Ethiopia. J. Health Pollution. 2021; 11(30):210601–210613.

[52] Mulati P, Kitur E, Taracha C, Kurgat J, Raina S, Irungu J. Evaluation of neonicotinoid residues in hive products from selected counties in Kenya. J. Environ. Anal. Toxicol. 2018; 8(4):1–7. 10.4172/2161-0525.1000577.

[53] Marete GM, Lalah JO, Mputhia J, Wekesa VW. Pesticide usage practices as sources of occupational exposure and health impacts on horticultural farmers in Meru County, Kenya. Heliyon. 2021;7(2):e06118.10.1016/j.heliyon.2021.e06118.

[54] Lalah JO, Otieno PO, Odira Z, Ogunah JA. Pesticides: Chemistry, manufacturing, regulation, usage and impacts on population in Kenya. Intech Open Journal. 2022.

[55] Boateng KO, Dankyi E, Amponsah IK, Awudzi GK, Amponsah E, Darko G. Knowledge, perception, and pesticide application practices among smallholder cocoa farmers in four Ghanaian cocoa-growing regions. Toxicology Reports. 2023;10:46–55. 10.1016/j.toxrep.2022.12.008

[56] Johannsmeier MF. Beekeeping in South Africa. Agricultural Research Council of South Africa, Plant Protection Research Institute; 2001.

[57] Masehela TS. An assessment of different beekeeping practices in South Africa based on their needs bee (forage use), services (pollination services) and threats (hive theft and vandalism). Doctoral dissertation, Stellenbosch: Stellenbosch University. 2017; pp. 239. 10.13140/RG.2.2.17985.66400.

[58] Wilson Rankin EE, Barney SK, Lozano GE. Reduced water negatively impacts social bee survival and productivity via shifts in floral nutrition. J. Insect Sci. 2020;20(5):1–8. 10.1093/jisesa/ieaa114.

[59] Squire DT, Richardson D, Risbey JS, Black AS, Kitsios V, Matear RJ, et al. Likelihood of unprecedented drought and fire weather during Australia’s 2019 megafires. npj Clim. Atmos. Sci. 2021;4(1): 1–12. 10.1038/s41612-021-00220-8.

[60] Tang KH, Yap PS. A systematic review of slash-and-burn agriculture as an obstacle to future-proofing climate change. In the Proceedings of the International Conference on Climate Change. 2020;4(1):1–19. 10.17501/2513258x.2020.4101

[61] Roe GH, Baker MB. Why is climate sensitivity so unpredictable? Science. 2007;318(5850):170–171.

[62] Li Y, Lu H, Yang K, Wang W, Tang Q, Khem S, Yang F, Huang Y. Meteorological and hydrological droughts in Mekong River Basin and surrounding areas under climate change. J. Hydrol. Reg. Stud. 2021;36:100873–100895. 10.1016/j.ejrh.2021.100873.

[63] Chemurot M, Kasangaki P, Francis O, Sande E, Isabirye-Basuta G. Beehive and honey losses caused by bush burning in Adjumani district, Uganda. Bee World. 2013;90(2):33–35. 10.1080/0005772x.2013.11417529.

[64] Winston ML, Otis GW, Taylor Jr OR. Absconding behaviour of the Africanized honeybee in South America. J. Apic. Res. 1979;18(2):85–94.

[65] F. Ruttner, “Biogeography and Taxonomy of Honey Bees.” 1988.

[66] Kipyatkov VE, editor. Life Cycles in Social Insects: Behaviour, Ecology and Evolution. St. Petersburg University Press; 2006.

[67] Schneider SS, McNally LC. Factors influencing seasonal absconding in colonies of the African honey bee, *Apis mellifera scutellata*. Insectes Soc. 1992;39(4):403–423. 10.1007/BF01240624.

[68] Kwadha CA, Ong’amo GO, Ndegwa PN, Raina SK, Fombong AT. The biology and control of the greater wax moth, *Galleria mellonella*. Insects. 2017;8(2):1–17.doi: 10.3390/insects8020061.

[69] Kuboja N, Kilima F, Isinika A. absconding of honeybee colonies from beehives: underlying factors and its financial implications for beekeepers in Tanzania. Int. J. Agric. Sci. Res. Technol. Ext. Educ. Syst. 2020;10(4): 85–193.

[70] Neumann P, Pirk C, Hepburn H, Solbrig A, Ratnieks F, Elzen P, Baxter J. Social encapsulation of beetle parasites by Cape honey bee colonies (*Apis mellifera capensis* Esch.). Naturwissenschaften. 2001;88(5):214–216. 10.1007/s001140100224.

[71] Wambua B, Muli E, Kilonzo J, Ng’ang’a J, Kanui T, Muli B. Large hive beetles: an emerging serious honey bee pest in the Coastal highlands of Kenya. Bee World. 2019;96(3):90–91. 10.1080/0005772x.2019.1568355.

[72] Payne AN, Shepherd TF, Rangel J.The detection of honey bee (*Apis mellifera*)-associated viruses in ants. Sci. Rep. 2020; 10(1):2923–2931. 10.1038/s41598-020-59712-x.

[73] Ord TJ. Drought-induced relocation of ant colonies and its consequences for the long-term spatial ecology of a population under stress. Funct. Ecol. 2023;37(8):2231–2245. 10.1111/1365-2435.14383.

[74] Spiewok S, Neumann P, Hepburn HR. Preparation for disturbance-induced absconding of Cape honeybee colonies (*Apis mellifera capensis* Esch.). Insectes Soc. 2006; 53(1): 27–31. 10.1007/s00040-005-0829-6.

[75] Guesh G, Amssalu B, Hailu M, Yayeneshet T. Epidemiology of honey bee disease and pests in selected zones of Tigray region, Northern Ethiopia. MSc. thesis, Bahir Dar University. 2015.

[76] Solomon S, Degu T, Fesseha H, Mathewos M. Study on major parasitic diseases of adult honeybees in three districts of Kaffa Zone, Southern Ethiopia. Vet. Med. Int. 2021; 2021:1–7. 10.1155/2021/6346703.

[77] Robi DT, Temteme S, Aleme M, Bogale A, Bezabeh A, Mendesil E. Health status of honey bee colonies (*Apis mellifera*) and disease-associated risk factors in different agroecological zones of Southwest Ethiopia. Vet. Parasitol. Reg. Stud. Reports. 2024;100943.

[78] Chemurot M, Akol AM, Masembe C, De Smet L, Descamps T, de Graaf DC. Factors influencing the prevalence and infestation levels of *Varroa destructor* in honey bee colonies in two highland agro-ecological zones of Uganda. Exp. Appl. Acarol. 2016; 68:497–508. 10.1007/s10493-016-0013-x.

[79] Boecking O, Ritter W. Grooming and removal behaviour of *Apis mellifera intermissa* in Tunisia against *Varroa jacobsoni*. J. Apic. Res. 1993;24(6):127–134. 10.1080/00033799300200371.

[80] Oldroyd BP, Allsopp MH. Risk assessment for large African hive beetles (*Oplostomus spp.*)—a review. Apidologie. 2017;48(4):495–503. 10.1007/s13592-017-0493-7.

[81] Meixner MD. A historical review of managed honey bee populations in Europe and the United States and the factors that may affect them. J. Invertebr. Pathol. 2010;103:S80–S95. 10.1016/j.jip.2009.06.011.

[82] Oberreiter H, Brodschneider R. Austrian COLOSS survey of honey bee colony winter losses 2018/19 and analysis of hive management practices. Diversity. 2020;12(3):99–126. 10.1016/j.jip.2009.06.011.

[83] Wheeler MM, Robinson GE. Diet-dependent gene expression in honey bees: honey vs. sucrose or high fructose corn syrup. Sci. Rep. 2014; 4(1): 5726, 2014, doi: 10.1038/srep05726.

[84] Fine JD, Shpigler HY, Ray AM, Beach NJ, Sankey AL, Cash-Ahmed A, et al. Quantifying the effects of pollen nutrition on honey bee queen egg laying with a new laboratory system. PloS one. 2018;13(9):e0203444. 10.1371/journal.pone.0203444

[85] Hoover SE, Ovinge LP, Kearns JD. Consumption of supplemental spring protein feeds by western honey bee (Hymenoptera: Apidae) colonies: effects on colony growth and pollination potential. J. Econ. Entomol. 2022; 115(2):417–429. 10.1093/jee/toac006.

[86] Topal E, Mărgăoan R, Bay V, Takma Ç, Yücel B, Oskay D, et al. The effect of supplementary feeding with different pollens in autumn on colony development under natural environment and in vitro lifespan of honey bees. Insects. 2022; 13(7):588–601. 10.3390/insects13070588.

[87] Pirk CW, Boodhoo C, Human H, Nicolson SW. The importance of protein type and protein to carbohydrate ratio for survival and ovarian activation of caged honeybees (*Apis mellifera scutellata*). Apidologie. 2010;;41(1):62–72. doi: 10.1051/apido/2009055.

[88] Altaye SZ, Pirk CW, Crewe RM, Nicolson SW. Convergence of carbohydrate-biased intake targets in caged worker honeybees fed different protein sources. J. Exp. Biol. 2010;213(19):3311–3318. doi: 10.1051/apido/2009055.

[89] Cunningham MM, Tran L, McKee CG, Polo RO, Newman T, Lansing L, et al. Honey bees as biomonitors of environmental contaminants, pathogens, and climate change. Ecol. Indic. 2022;134:108457–1088467. doi: 10.1051/apido/2009055.

[90] T. J. Meutchieye F., Ngamadjeu D.D, “Beekeeping features in Cameroon Adamawa grasslands Beekeeping features in the Cameroon Adamawa grasslands,” Genet. Biodivers. J., vol. 2, pp. 11–16, 2018.

